# Come together: bioelectric healing-on-a-chip

**DOI:** 10.1101/2020.12.29.424578

**Authors:** Tom J. Zajdel, Gawoon Shim, Daniel J. Cohen

**Affiliations:** Department of Mechanical & Aerospace Engineering, Princeton University, Princeton, NJ, United States of America

## Abstract

There is a growing interest in bioelectric wound treatment and electrotaxis, the process by which cells detect an electric field and orient their migration along its direction, has emerged as a potential cornerstone of the endogenous wound healing response. Despite recognition of the importance of electrotaxis in wound healing, no experimental system to date demonstrates that the actual closing of a wound can be accelerated solely by the electrotaxis response itself, and in vivo systems are too complex to resolve cell migration from other healing stages such as proliferation and inflammation. This uncertainty has led to a lack of standardization between stimulation methods, model systems, and electrode technology required for device development. In this paper, we present a ‘healing-on-chip’ approach that is a standardized, low-cost, model for investigating electrically accelerated wound healing. Our device provides the first convergent field geometry used in a stimulation device. We validate this device by using electrical stimulation to close a 1.5 mm gap between two large (30 mm^2^) primary skin keratinocyte layers to double the rate of healing over an unstimulated tissue. This proves that convergent electrotaxis is both possible and can accelerate healing, and offers a new ‘healing-on-a-chip’ platform to explore future bioelectric interfaces.

## Introduction

Since du Bois-Reymond first characterized the naturally occurring ‘wound current’ nearly two centuries ago (1), there has been significant interest in applying external electrical stimulation to improve wound healing (2–4). The potential for this approach is becoming increasingly apparent— for instance, numerous, recent, *in vivo* studies show some improvement in skin healing in animal models upon electric field stimulation (5–10), while *in vitro* assays have demonstrated control of cells and simple tissues using spatially programmed electric cues (11–13). Further, given the increasing prevalence and healthcare burden of wound treatments (14,15), new technologies to expedite and improve wound care are sorely needed. However, despite these and other studies over the past several decades, the few extant commercial products have demonstrated mixed results (16–19), and bioelectric wound therapy is far from the standard of care. This discrepancy is due to broad gaps in both technology development and biological knowledge describing how electrical stimulation may act to improve wound healing. Technologically, optimum stimulation parameters for field strength, biointerface design, and current delivery mode remain unclear (20,21). Biologically, there is uncertainty about how the key wound healing mechanisms--cell migration, proliferation, and inflammation—are affected by electric stimulation (2). This uncertainty has resulted in a lack of standardization in stimulation schemes, model systems, and technology that can all lead to issues of reproducibility and long design iterations time that have slowed progress (22–24).

Here, we begin to address this problem by integrating a popular technical approach used in other branches of biotechnology—’organ-on-a-chip’ systems—to reduce the complexity of biomedical problems to something both tractable and eventually translatable. Organ-on-a-chip (OoC) platforms are *in vitro* model systems that capture a specific and critical physiological behavior of the *in vivo* system in a standardized, rapid, lower-cost *in vitro* model. To date, OoCs have clearly proven their value in other fields by aiding discoveries and treatments for lung, gut, and vascular pathologies (25–27). Here, we use an OoC approach to integrate a ‘healing-on-a-chip’ platform with a custom electrobioreactor designed from the ground up to investigate electrically accelerated wound healing.

While there are many effects that applied electrical stimulation may have on tissue growth and healing, the best-characterized is *electrotaxis*—the directed motion of cells in response to an electric current. Electrotaxis is seen in over 20 cell types across multiple organisms where cells sense and track electrochemical potential gradients (∼1 V/cm) that emerge during development and injury healing (28–30). The mechanism of detection is thought to be electrophoresis of charged membrane-bound receptors in the presence of an electric field, resulting in an asymmetric distribution of these proteins that triggers downstream signaling of the cell migration machinery (31). *In vivo*, these fields result in the center of a skin wound being negatively polarized relative to the periphery of the wound (32,33). Direct current fields are analogous to fields *in vivo* (34), in contrast to the pulsed DC or AC stimulation used in many *in vivo* studies (5), and are sufficient to induce the electrotaxis response. However, electrotaxis has primarily been studied in isolated single cells to elucidate the molecular biology of the process, and at present there is no study, either *in vitro* or *in vivo* that conclusively indicates that electrotaxis itself can accelerate wound closure. This gap stems from the technological limitations of current devices used to study electrotaxis.

Nearly all devices use a single electrode pair to apply a uniform, unidirectional field across tissues—such a field would cause one side of a wound to close and the other side to worsen. *In vivo*, the wound field converges on the center of a wound, so a new device design is required to capture this characteristic. In addition to this stimulation limitation, most studies and devices do not generalize well to macroscale tissues and wounds since precise tissues with reproducible, millimetric wounds must be grown inside the electrobioreactor. Finally, macroscale cell migration requires stable electrical stimulation over many hours, and the common, bleach-based electrode preparation process is insufficient for long-term stimulation (>4 hours). An ideal bioelectric ‘wound-on-a-chip’ platform should address these issues.

Here we build on our prior work (11,12) to create a new electrobioreactor to study healing in a macroscale skin-on-a-chip model using primary mouse skin monolayers which migrate toward the cathode when stimulated (Fig. 1), and use electrical stimulation to accelerate closure of 1.5 mm large model skin wounds by at least 2X over unstimulated skin layers (Figs. 2, 3). To accomplish this, we developed new electrotaxis infrastructure specifically designed for the constraints of wound healing, delivering a sustained converging electric field to a tissue (Fig. 1). With this device, we were able to engineer and stimulate the largest tissues yet tested with electrotaxis (30 mm^2^) for 12 hours, while also exploring the consequences of overstimulation.

**Fig. 1:**
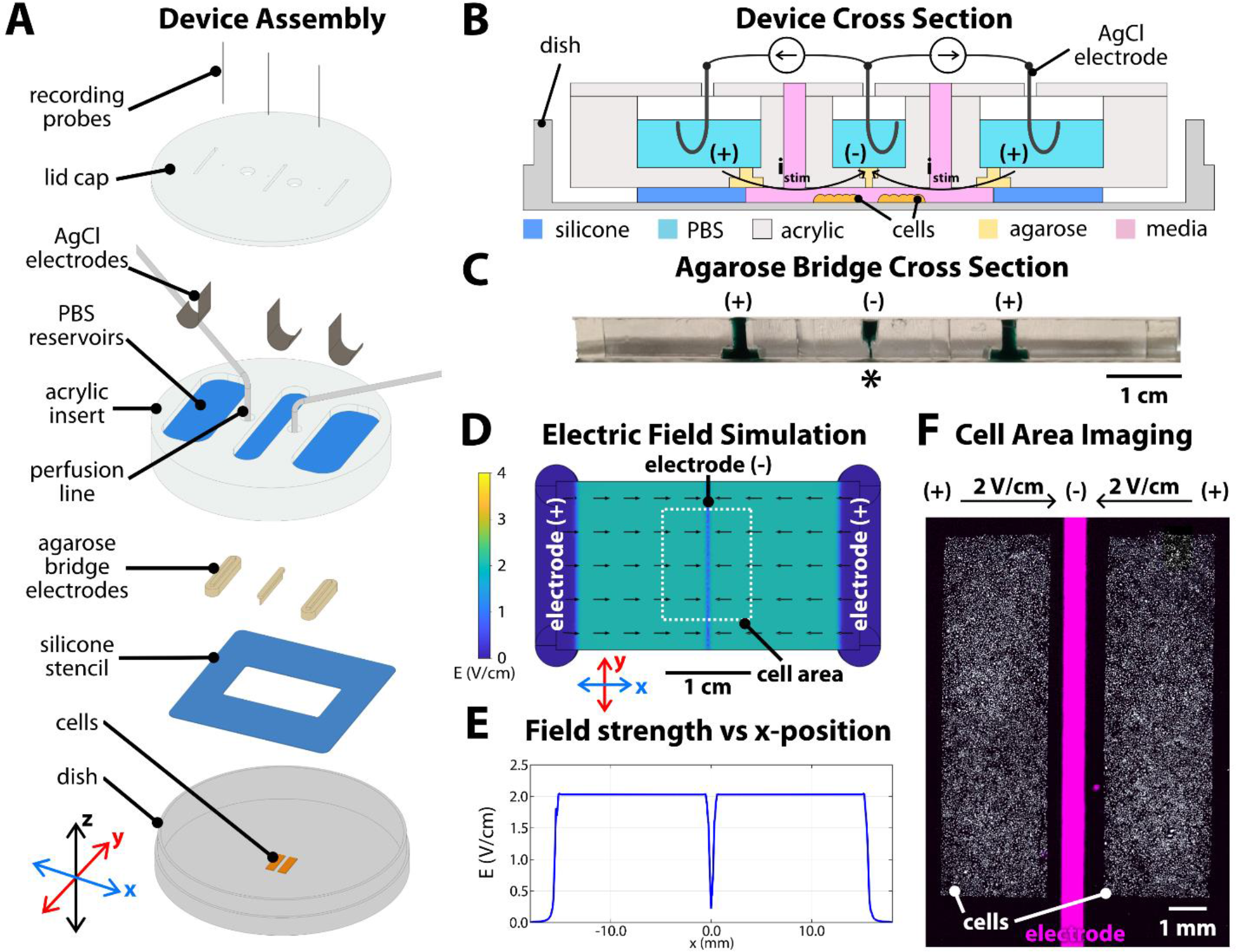
Convergent field stimulation device. (A) Layer-based assembly of the bioreactor onto a tissue culture dish. Cells are patterned in the center of the dish, then a 250 µm-thick silicone stencil is placed to define the stimulation area and height. Agarose bridges are cast inside an acrylic insert, then clamped into the dish and against the silicone stencil. The reservoirs on the topside of the acrylic insert are filled with phosphate-buffered saline (PBS). Chloridized silver electrodes and titanium wire recording probes are inserted in each reservoir, all held in place by a lid cap. (B) Device cross-section sketch and (C) photograph of the sectioned agarose bridges stained with green food coloring for contrast. The narrow cathode is labeled with ‘*’. (D) Numeric simulation of the electric field in the device, showing constant 2 V/cm field strength converging toward the center, with a steep drop-off in strength starting ±500 µm from the center. (E) Simulated field strength versus x-position in the device. (F) Microscope capture of the central area of the assembled device, showing the central electrode 500 µm wide positioned between the two tissues. The cells (white) were labeled with a Cy5 lipophilic dye and the outline of the central electrode was visualized with a DAPI filter set (λ_ex_/ λ_em_ 358/461 nm) and filled via post-processing in ImageJ (magenta).

**Fig. 2.**
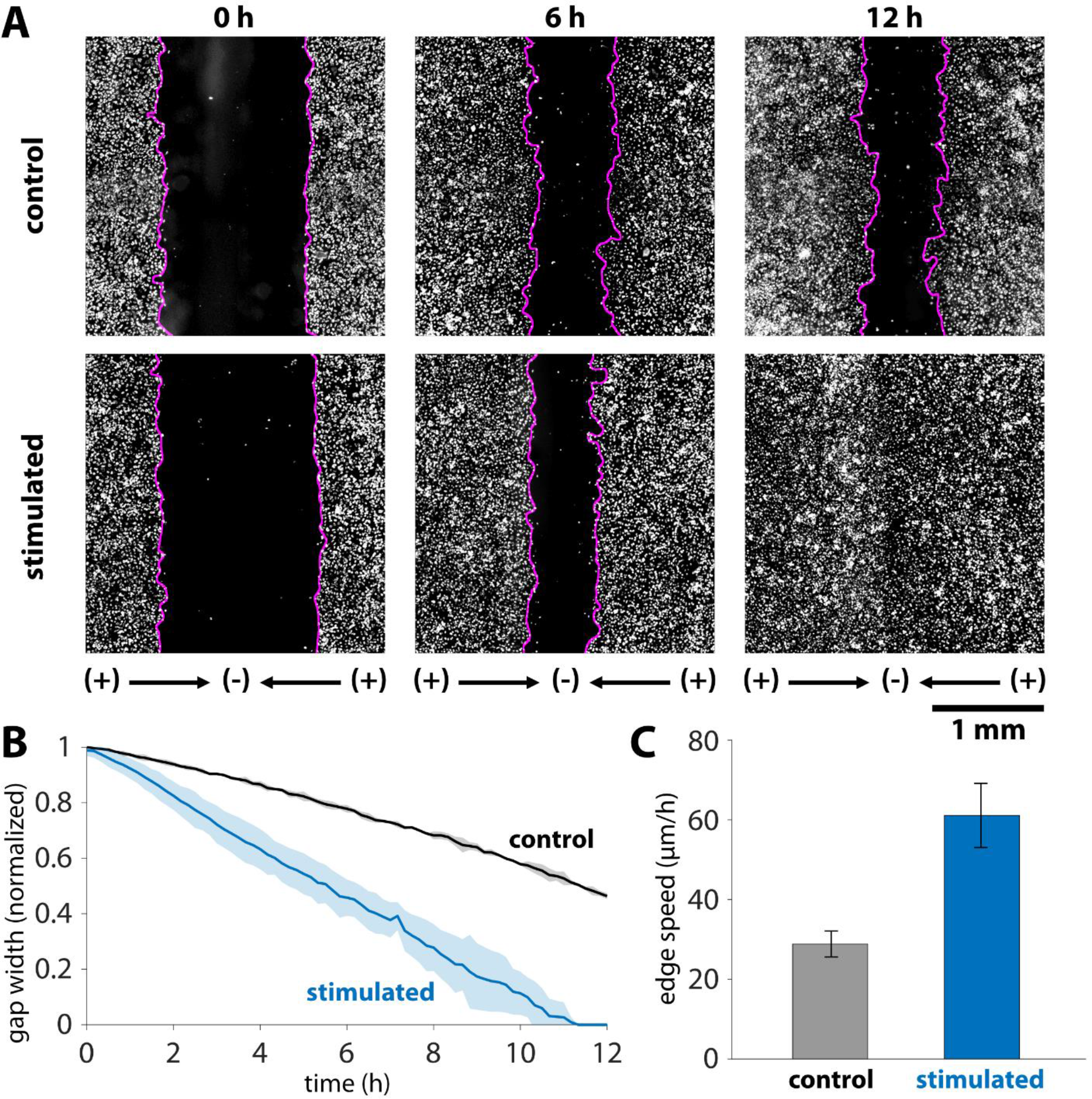
Dynamics of accelerated wound closure of keratinocyte monolayers. (A) Timepoint comparison of stimulation versus control for keratinocytes labeled with a Cy5 cytoplasmic dye. Gap boundaries are demarked by magenta lines. Initial gap between tissues was 1.5 mm, and this gap closed by 12 hours in the stimulated case, while roughly 50% of the gap remained in the control case. (B) Gap closure normalized to the initial gap width for N = 3 tissues in each condition. Shaded region represents standard deviation. (C) Edge expansion speeds averaged over an 8 h period. The average edge expansion was 29.4 ± 3.3 µm/h and 62.2 4 ± 8.1 µm/h for the control and stimulated cases, respectively. N = 6 edges in each case and error bars represent standard deviation.

**Fig. 3.**
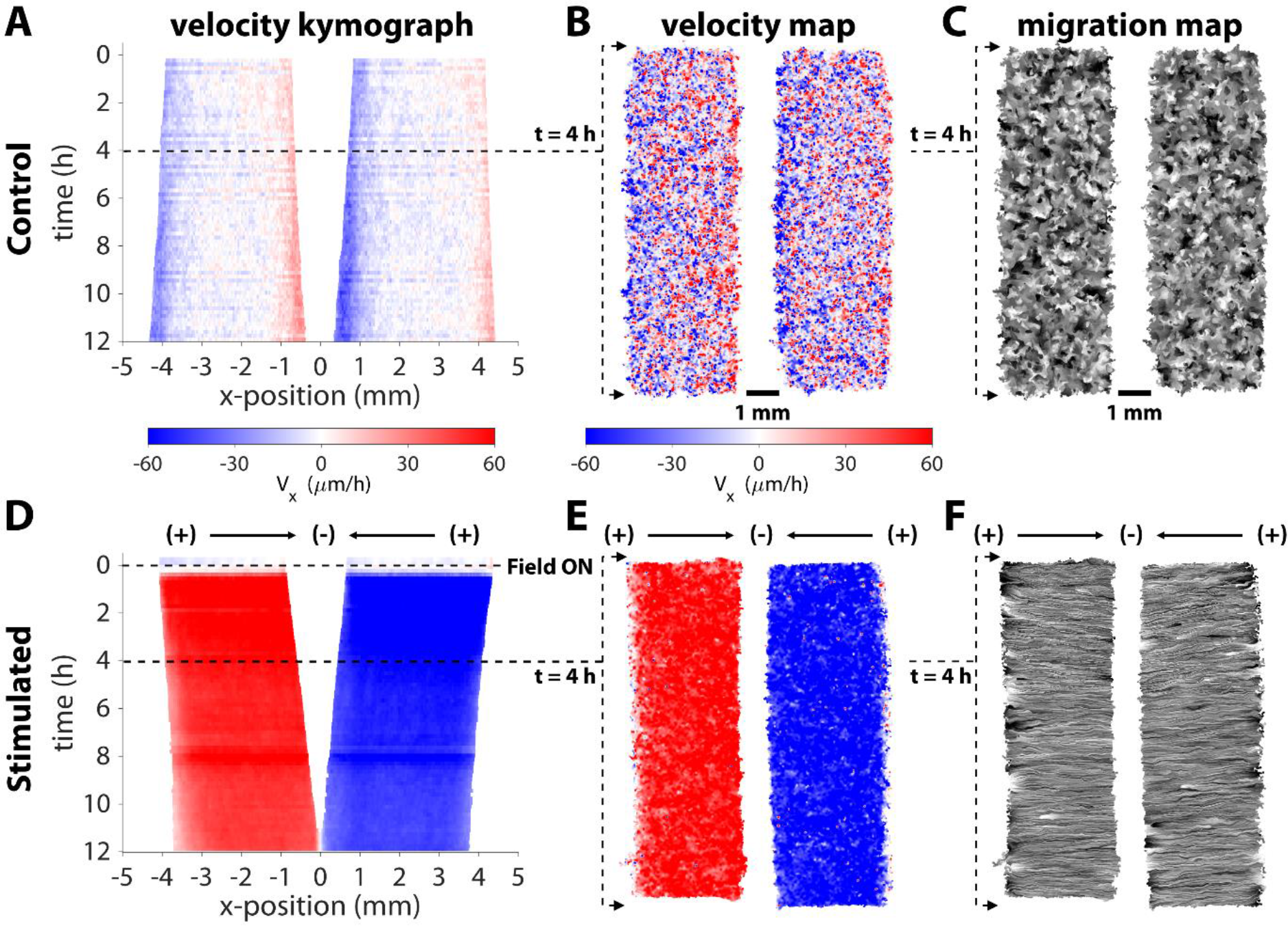
Convergent stimulation results in coherent migration response towards the center. Representative kymograph of Vx averaged across x-position in a control (A) and stimulated (D) tissue pair. The control tissues expand uniformly outwards, while the stimulated tissues converge towards the cathode with uniformly high speed across the tissue area (∼60 µm/h directed towards the gap). Horizontal velocity maps (B,E) and migration maps (C,F) in representative tissue pairs.

## Results

Our new electrobioreactor significantly departs from extant electrotaxis systems by generating an electric field that converges at the center of model wounds, and functions as follows. The device consists of an acrylic insert clamped to a standard tissue culture dish, holding electrodes and agarose salt bridges in position (Fig. 1A, see ESI Methods for fabrication details). Three chloridized silver stimulation electrodes (anodes at left and right, cathode at center) are isolated from each other in separate saline reservoirs and electrical contact with the culture media is provided by 4% agarose w/v salt bridges cast inside the insert, one per reservoir (Fig. 1B). The ∼500 µm thin, laser-milled, agarose bridge serves as a central cathode and is aligned directly over the wound site (Fig. 1C, see ‘*’). The result is a stable, uniform field that converges upon the central electrode as confirmed by simulation (Fig. 1D,E). To reliably generate reproducible tissues and linear wounds, we use a silicone stencil templating method (12,35) to prepare confluent monolayers the evening before an experiment. We then assemble the electrobioreactor over these tissues prior to imaging. For these experiments, we use layers of keratinocytes from primary skin cultured under basal conditions optimized for electrotaxis (11). After the tissues have grown, the stencils are removed, and the device is clamped over the cells, aligned such that that the central slit electrode is in the gap between cells (Fig. 1E). Then, a computer-controlled source meter (Keithley 2450) is connected to each pair of electrodes (left-center and right-center), with both sources sharing the central cathode, to supply an electric current. The field within the chamber is continuously monitored by a digital oscilloscope and the current output of each source meter is adjusted via closed-loop control to maintain a constant 2 V/cm field strength directed toward the central cathode. We specifically chose this field strength as it has previously been validated and was chosen to amplify the approximate field strength experienced *in vivo* (28). To extend cathode lifetime, only one source was active at a time, alternating between left-center and right-center stimulation every 30 seconds. Oxygen delivery and waste management are handled by perfusing fresh media through the bioreactor at 2 mL/h, turning over the chamber volume ∼11 times per hour. The resulting system provides a robust convergent field to viable cells.

Stable DC stimulation and cell viability require that the electrodes remain intact throughout an entire experiment, so optimization of electrode chemistry is an important consideration. Virtually all DC electrotaxis chambers use an anode and a cathode to inject Faradaic current through a sample, using combinations of salt bridges, media perfusion, and heavy buffering to prevent the buildup of toxic electrochemical byproducts or harmful pH changes due to electrolysis at the electrodes (20,36,37). Because the current used in our device is moderate (∼6-10 mA) and the 1.5 mm gap between tissues is relatively large, the central cathode must be able to sink current for an extended period to induce tissue convergence, ideally 12 hours or more. To support this, our system uses electrically chloridized silver foil as electrodes, which degrades at the cathode into ionic silver and chloride during stimulation. This reaction is more favorable than the hydrolysis cathodal half-reaction, which evolves hydrogen gas from the solution and increases pH (37). This allows for safe stimulation until AgCl is depleted at the cathode, when evolution of H_2_ then becomes favorable and pH increases rapidly, which can cause cytotoxicity. Therefore, sufficient chloridization of the silver foil is paramount for extended electrode lifetime. We compared our chloridization method with bleach immersion, another technique commonly used to chloridize silver. We performed repeated cyclic voltammetry to compare electrode preparations and found that our method of electroplating silver chloride resulted in more stable cathodes (see ESI, Fig. S1). Our combined approach of robust silver chloridization, agarose diffusion barriers to prevent ionic silver reaching the tissue, and media perfusion integrates numerous best practices to maximize cell viability during stimulation in our device, allowing for extended wound healing experiments.

To evaluate this platform for *in vitro* healing, we patterned two 10 x 3 mm tissues spaced 1.5 mm apart with the central cathode aligned over the wound center (Fig. 1F). The acrylic outline of the central cathode slit fluoresces weakly when imaged using a standard DAPI filter set, so the alignment between the central cathode and the tissues could be tuned and verified. We then applied convergent electrical stimulation over 12 hours following a 30-minute control period without field, with striking results (Fig. 2, Video S1). In the non-stimulated control case, cell proliferation and migration lead to the slow expansion of tissues and gradual, but incomplete closure of the wound over 12 hours (∼50% closure, N=3). However, convergent bioelectric stimulation led to complete closure between 11-12 h (N=3). More specifically, the edge migration speed was twice as fast in the stimulated case as in the control, measuring 29.4 ± 3.3 µm/h and 62.2 ± 8.1 µm/h for the control and stimulated cases, respectively. To conclusively attribute this effect to electrical stimulation rather than temperature effects (Joule heating has been linked to increased migration speeds in prior studies (31,38)) we monitored the device temperature during stimulation (Fig. S2). The steady state temperature rose from 37 °C to 38 °C, and this 3% increase is unlikely to account for the 100+% increase in migration speed during stimulation. We hypothesize that perfusion and media turn over helps to exchange heat and mitigate any effects from Joule heating. Taken together, this is the first demonstration of convergent field stimulation accelerating *in vitro* wound healing, and the results prove that electrotaxis alone is sufficient for this closure.

To better characterize device performance and its effects on large scale tissue growth and motion, we performed particle image velocimetry (PIV) on each tissue. Representative horizontal velocity kymographs for both the control and convergent stimulation cases are shown in Fig. 3 (compare with Video S1). To provide context of spatial dynamics within a given tissue, we show representative heatmaps of horizontal velocity and line integral convolution (LIC) migration maps to visualize the overall flow of cellular motion at 4 hours after the onset of stimulation (steady state). Throughout the control tissue (Figs. 3A-C), there is little net outwards motion, except for slow expansion at the edges. Disorder is apparent in the velocity and migration maps of the control tissues, which lack large regions of coordinated movement, as expected for non-stimulated tissues (Figs. 3B,C). In contrast, bioelectric stimulation resulted in nearly uniformly high-speed motion throughout the tissue, converging on the gap within 15 minutes of the field turning on, as visualized in the velocity and migration maps (Figs. 3D-F).

The large number of parallel streaklines along the stimulation direction in the migration map demonstrates highly coordinated motion across the tissue in alignment with the stimulus (Figs. 3F). These visualizations reveal that the electric field acts a global migration cue across a large area, confirming that cells experience a highly uniform field as predicted by simulation (Fig. 1D,E).

Having demonstrated that the *in vitro* healing process can be electrically accelerated overall, we next characterized cellular responses specifically during the final stages of wound closure. Unlike traditional electrotaxis chambers where the electrodes are significantly distal to the tissue to ensure a uniform field, our healing-on-a-chip device requires a central electrode to focus cell migration into the wound zone. Since the central electrode has a finite width (∼500 µm here) that is smaller than the wound, this means that tissues will eventually pass underneath the electrode and enter the ‘electrode shadow’ during the final stages of healing and convergence. Any discrete electrode produces field non-uniformities close to its surface, so as cells enter the electrode shadow, they will experience a very different field than out in the fully developed zones far from the center. Our simulation predicts a sharp decrease in electric field strength that begins about 500 µm on either side of the central cathode above the convergence region (Fig. 4A). We quantified the actual effects of the central field singularity by stimulating closed tissues for 6 hours and using a live nuclear dye to track cells in that central zone (Video S2). We averaged PIV across the region surrounding the closure zone over the stimulation period (Fig. 4B, asterisks and error bars) and fit a sigmoid function to the data (Fig. 4B, inset) showing that there is a strong, steady-state response far from the central electrode that steadily weakens as cells approach the central electrode and enter the electrode shadow (Fig. 4, dashed blue line; magenta zone shows electrode shadow). While this local weakening of the electrotactic response closely resembled the trend in our simulations, cells nonetheless continued to directionally migrate deep into the electrode shadow zone, only to dropping to <50% of the steady state velocity once cells were ∼100 µm off the electrode midline. These data show that the effective electrode size is smaller than its physical, 500 µm width (Fig. 4B, compare dotted black boundaries to electrode boundaries). This means that even relatively large electrodes can still promote last-mile healing.

**Fig. 4.**
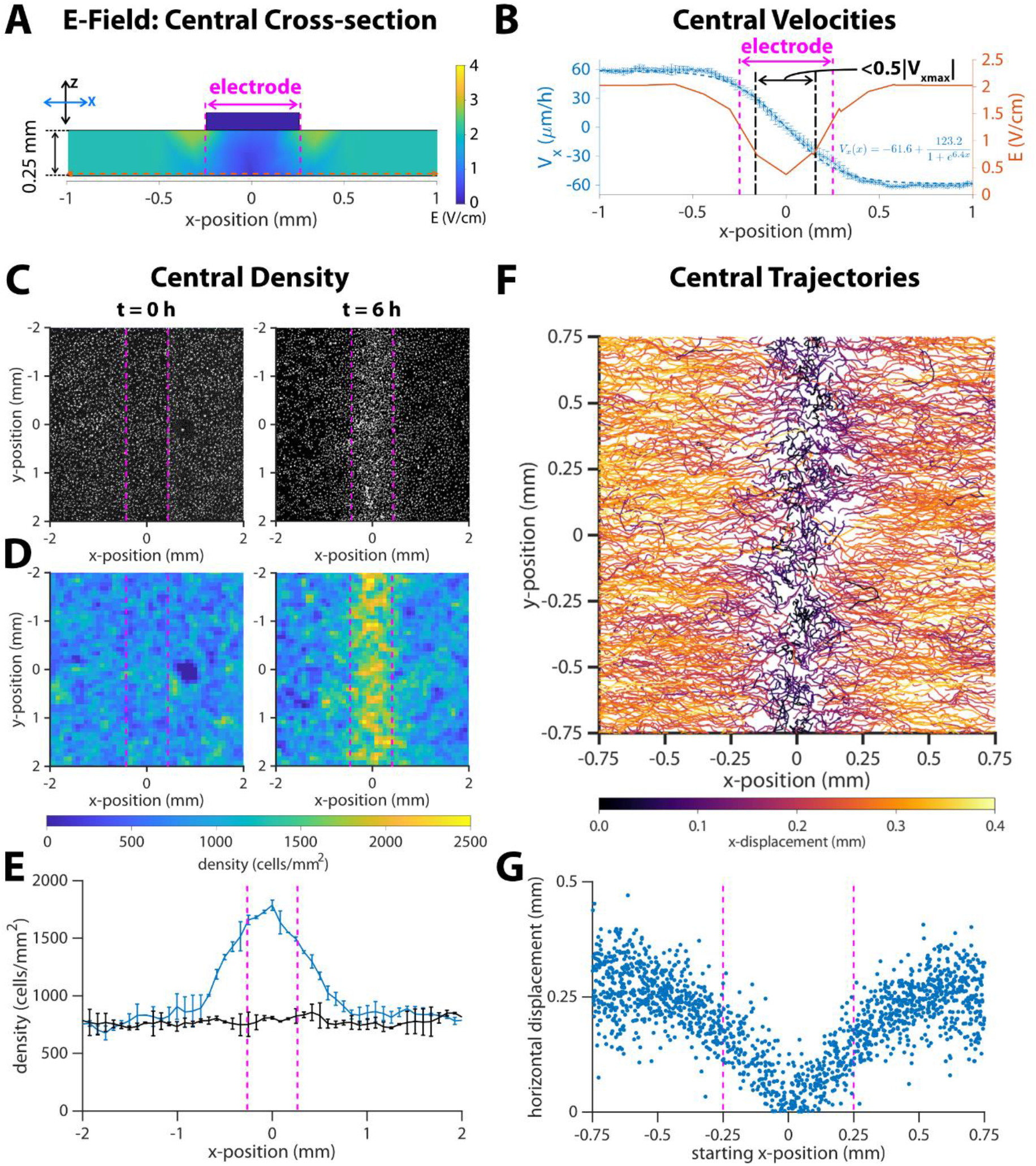
Response of migrating cells near the center of a closed tissue. Dotted vertical magenta lines indicate approximate cathode boundaries in each respective graph. (A) Numeric simulation results for the electric field of the channel in cross section. The horizontal dotted line is the section of the electric field plotted in the next panel. (B) Average horizontal velocity plotted as blue *’s with error bars representing standard deviation (N = 2). The dashed blue line indicates a least-squares fit of a sigmoid function to the data, and the formula for this fit is inset in the lower right quadrant of the plot. The region where the horizontal velocity’s magnitude drops to < 50% of the steady state value is marked by dot-dash black lines. Predicted strength from numeric simulation (reproduced from Fig. 1E) is plotted as a red solid line. (C) DAPI images of cell nuclei and (D) density maps for tissues at the onset of stimulation (t = 0 h) and after six hours of stimulation (t = 6 h). (E) Average cell density versus x-position, where x = 0 is the center of the cathode. Error bars represent standard deviation (N = 2). (F) Montage of 6-hour trajectories of individual cells in proximity with the center. Each track is colored by net x-displacement. (G) Total horizontal displacement of cell versus its starting x-position relative to the center of the cathode at x = 0.

Critically, we also observed potential consequences to continued electrical stimulation after a wound had closed. As has been noted previously (39), electrotaxis appears to override basic cellular safety mechanisms, such as contact inhibition, meaning that cells will continue trying to directionally migrate as long as stimulation is active. In our wound healing model, this meant that stimulating after a tissue had closed would continue to drive cell migration towards where the center of the wound had previously been. This inevitably caused an increase in local cell density, and we measured a >2X increase (from 750 to 1600 cells/mm^2^) in cell density under the electrode shadow relative to density distal to the central electrode (Fig. 4C-E), showing there is potential for over-densification of cells driven by post-healing stimulation. Comparing individual cell trajectories within the central zone confirmed that cells within the electrode shadow translated horizontally a lower distance than those that were farther away (Fig. 4F,G). This reduction in overall translation extended, in a graded fashion, outwards 500 µm from the center in either direction, consistent with the reduction of speed cells experience as they enter the electrode shadow. Nevertheless, there is net migration towards the center, even for cells that were initially positioned under the electrode, suggesting again that the electrode’s influence extends underneath its width despite significant weakening of the effective field strength.

We suggest two reasons that the ‘effective electrode’ size would be smaller than its physical size. First, the threshold field strength that elicits an electrotaxis response is lower than the 2 V/cm we target in stimulation. As the field strength rolls off, it is still ‘therapeutic’ for some time, given that physiological field strengths are on the order of 1 V/cm (28). Second, the monolayers carry some memory of the electrotaxis response that continues to influence their responses after the stimulus changes (11). Keratinocytes polarize in response to the field stimulus, and this polarization takes time to decay once a stimulus is no longer detected. This could lead to cells effectively coasting, unguided, during the last gap before tissue closure.

## Discussion & Conclusion

Overall, we present a bioelectric, healing-on-a-chip (HoC) platform designed specifically to study the role of electrotaxis and other electrical phenomena in wound healing. Unique for electrical stimulation bioreactors, our approach creates a field stimulation pattern that mimics that found in wounds *in vivo*, with the field converging at the center of the wound gap. This capability allows us to directly explore the actual healing process, rather than purely uni-directional cell migration. Therefore, our platform allows study of the *in vitro* healing process spanning initial injury, ‘first contact’ as the sides of the wound meet and, critically, post-closure behavior after the wound has healed. Using this platform and unoptimized stimulation parameters, we demonstrate ∼2X acceleration of wound closure in an *in vitro* skin layer model due solely to electrotactic effects. To our knowledge, this is the first demonstration and visualization of electrotaxis itself accelerating a healing process. The stability, reproducibility, and programmability of the platform make it suitable to deeply explore key technological and biological questions, and we have taken care to ensure the device is easily replicable and accessible to a broad audience.

That even naïve stimulation had a strong, positive effect on *in vitro* healing is encouraging, and establishes a clear baseline against which future parameter optimization studies can be compared. This approach could be critical to the field as standardization and optimization of stimulation approaches remains an open question. To hint at this, we explored the effects of stimulating beyond initial closure of the wound by electrically stimulating a closed tissue, and the resulting cellular pile-up indicates both the potency of electrotaxis to drive migration and the importance of being able to fine-tune and intelligently adjust stimulation in practice to avoid detrimental effects of overstimulation. Such cellular pile-ups also speak more fundamentally to the role of electrotaxis as a tool to modulate and explore interactions at the boundaries between tissues.

While we specifically investigated healing in monolayers of primary skin cells here, wound healing *in vivo* clearly involves complex coordination across multiple cell types (e.g. macrophages and immune cells, fibroblasts, and vascular cells, and epidermal cells) and phases, (e.g inflammation, granulation, and re-epithelialization) (2). That our platform supports pre-engineering tissue configurations means that co-cultures or more complex tissue models can be grown first and then incorporated into the bioreactor to allow more complex studies on healing. When linked to stimulation optimization approaches, it may be possible to determine modalities that preferentially target a given cell type, or process such as proliferation vs. migration during healing. Again, these questions benefit from a field geometry that enables a healing phenotype.

Finally, our bioelectric ‘Healing-on-a-Chip’ approach is fully open and intended to be modified and tailored for a variety of applications. We provide complete design files, computational models, and stimulation code (Provided via a GitHub repository: https://github.com/CohenLabPrinceton/SCHEEPDOG), and the basic approach lends itself to easy customization. For instance, electrode shape, size, number, and location can easily be adjusted without additional cost or significant complexity. Field stimulation strategies can be tested by attaching any desired power supplies or running arbitrary stimulation code to activate electrode sequences. Our autofluorescence alignment approach makes it possible to accurately align a given electrode configuration to a given wound and removes much of the ambiguity and difficulty this process would normally introduce. We hope the demonstrations here and flexibility of the device can help accelerate healing-on-a-chip research, improve translation for future *in vivo* applications, and even support new, research on general interactions between colliding tissues.

## Supporting information

Electronic Supplementary Information

## Acknowledgements

We gratefully acknowledge Prof. Danelle Devenport and Katie Little at Princeton University for providing primary keratinocytes and culture support. Research reported in this publication was supported by the National Center for Advancing Translational Sciences (NCATS), a component of the National Institute of Health (NIH) under award number TL1TR003019 (TJZ). Further support was provided by National Institutes of Health grant R35GM13357401 (DJC, GS). The content is solely the responsibility of the authors and does not necessarily represent the official views of the National Institutes of Health.

## Notes

### Competing Interest Statement

The authors have declared no competing interest.

## References

1. Bois-Reymond E du. Untersuchungen uber thierische Elektricitat. Berlin, Reimer. 1848;1.

2. Tai G, Tai M, Zhao M. Electrically stimulated cell migration and its contribution to wound healing. Burns & trauma. Narnia; 2018;6(1).

3. Anderson CA, Hare MA, Perdrizet GA. Wound healing devices brief vignettes. Advances in wound care. Mary Ann Liebert, Inc. 140 Huguenot Street, 3rd Floor New Rochelle, NY 10801 USA; 2016;5(4):185–190.

4. Hunckler J, De Mel A. A current affair: electrotherapy in wound healing. Journal of multidisciplinary healthcare. Dove Press; 2017;10:179.

5. Long Y, Wei H, Li J, Yao G, Yu B, Ni D, et al. Effective Wound Healing Enabled by Discrete Alternative Electric Fields from Wearable Nanogenerators. ACS Nano. 2018;12(12):12533–12540.

6. Liang Y, Tian H, Liu J, Lv YL, Wang Y, Zhang JP, et al. Application of stable continuous external electric field promotes wound healing in pig wound model. Bioelectrochemistry [Internet]. Elsevier LTD; 2020;135:107578. Available from: https://doi.org/10.1016/j.bioelechem.2020.107578

7. Jang HK, Oh JY, Jeong GJ, Lee TJ, Im GB, Lee JR, et al. A disposable photovoltaic patch controlling cellular microenvironment for wound healing. International Journal of Molecular Sciences. 2018;19(10).

8. Kai H, Yamauchi T, Ogawa Y, Tsubota A, Magome T, Miyake T, et al. Accelerated Wound Healing on Skin by Electrical Stimulation with a Bioelectric Plaster. Advanced Healthcare Materials. 2017;6(22):1–5.

9. Yu C, Xu ZX, Hao YH, Gao YB, Yao BW, Zhang J, et al. A novel microcurrent dressing for wound healing in a rat skin defect model. Military Medical Research [Internet]. Military Medical Research; 2019;6(1):1–9. Available from: https://mmrjournal.biomedcentral.com/track/pdf/10.1186/s40779-019-0213-x

10. Wang X-F, Li M-L, Fang Q-Q, Zhao W-Y, Lou D, Hu Y-Y, et al. Flexible electrical stimulation device with Chitosan-Vaseline® dressing accelerates wound healing in diabetes. Bioactive materials. Elsevier; 2020;6(1):230–243.

11. Zajdel TJ, Shim G, Wang L, Rossello-Martinez A, Cohen DJ. SCHEEPDOG: Programming Electric Cues to Dynamically Herd Large-Scale Cell Migration. Cell Systems [Internet]. Elsevier Inc.; 2020;10(6):506–514.e3. Available from: https://doi.org/10.1016/j.cels.2020.05.009

12. Cohen DJ, Nelson WJ, Maharbiz MM. Galvanotactic control of collective cell migration in epithelial monolayers. Nature materials. Nature Publishing Group; 2014;13(4):409–417.

13. Gokoffski KK, Jia X, Shvarts D, Xia G, Zhao M. Physiologic electrical fields direct retinal ganglion cell axon growth in vitro. Investigative ophthalmology & visual science. The Association for Research in Vision and Ophthalmology; 2019;60(10):3659–3668.

14. Padula WV, Delarmente BA. The national cost of hospital-acquired pressure injuries in the United States. International Wound Journal. 2019;16(3):634–640.

15. Martinengo L, Olsson M, Bajpai R, Soljak M, Upton Z, Schmidtchen A, et al. Prevalence of chronic wounds in the general population: systematic review and meta-analysis of observational studies. Annals of epidemiology. Elsevier; 2019;29:8–15.

16. Morris C. Bio-electrical stimulation therapy using POSiFECT®RD. Wounds UK. 2006;2(4):112–116.

17. Kloth LC. Electrical Stimulation Technologies for Wound Healing. Advances in Wound Care. 2014;3(2):81–90.

18. Kim H, Izadjoo M. Antibiofilm efficacy evaluation of a bioelectric dressing in mono-and multi-species biofilms. Journal of wound care. MA Healthcare London; 2015;24(Sup2):S10–S14.

19. Kim H, Park S, Housler G, Marcel V, Cross S, Izadjoo M. An overview of the efficacy of a next generation electroceutical wound care device. Military Medicine. Oxford University Press; 2016;181(Suppl_5):184–190.

20. Zhao Z, Zhu K, Li Y, Zhu Z, Pan L, Pan T, et al. Optimization of Electrical Stimulation for Safe and Effective Guidance of Human Cells. Bioelectricity. Mary Ann Liebert, Inc., publishers 140 Huguenot Street, 3rd Floor New …; 2020;

21. Khouri C, Kotzki S, Roustit M, Blaise S, Gueyffier F, Cracowski J-L. Hierarchical evaluation of electrical stimulation protocols for chronic wound healing: An effect size meta-analysis. Wound Repair and Regeneration. Wiley Online Library; 2017;25(5):883– 891.

22. Isseroff RR, Dahle SE. Electrical stimulation therapy and wound healing: where are we now? Advances in wound care. Mary Ann Liebert, Inc. 140 Huguenot Street, 3rd Floor New Rochelle, NY 10801 USA; 2012;1(6):238–243.

23. Kloth LC. Electrical stimulation for wound healing: a review of evidence from in vitro studies, animal experiments, and clinical trials. The international journal of lower extremity wounds. Sage Publications Sage CA: Thousand Oaks, CA; 2005;4(1):23–44.

24. Ashrafi M, Alonso-Rasgado T, Baguneid M, Bayat A. The efficacy of electrical stimulation in lower extremity cutaneous wound healing: a systematic review. Experimental dermatology. Wiley Online Library; 2017;26(2):171–178.

25. Zhang B, Korolj A, Lai BFL, Radisic M. Advances in organ-on-a-chip engineering. Nature Reviews Materials. Nature Publishing Group; 2018;3(8):257–278.

26. Wang Z, Samanipour R, Koo K, Kim K. Organ-on-a-chip platforms for drug delivery and cell characterization: A review. Sens. Mater. 2015;27(6):487–506.

27. Zhang B, Radisic M. Organ-on-a-chip devices advance to market. Lab on a Chip. Royal Society of Chemistry; 2017;17(14):2395–2420.

28. McCaig CD, Song B, Rajnicek AM. Electrical dimensions in cell science. Journal of cell science. The Company of Biologists Ltd; 2009;122(23):4267–4276.

29. Cortese B, Palama IE, D’Amone S, Gigli G. Influence of electrotaxis on cell behaviour. Integrative Biology. Oxford University Press; 2014;6(9):817–830.

30. Kennard AS, Theriot JA. Osmolarity-independent electrical cues guide rapid response to injury in zebrafish epidermis. Elife. eLife Sciences Publications Limited; 2020;9:e62386.

31. Allen GM, Mogilner A, Theriot JA. Electrophoresis of cellular membrane components creates the directional cue guiding keratocyte galvanotaxis. Current Biology. Elsevier; 2013;23(7):560–568.

32. Wahlsten O, Skiba JB, Makin IRS, Apell SP. Electrical field landscape of two electroceuticals. Journal of Electrical Bioimpedance. 2016;7(1):13–19.

33. Shen Y, Pfluger T, Ferreira F, Liang J, Navedo MF, Zeng Q, et al. Diabetic cornea wounds produce significantly weaker electric signals that may contribute to impaired healing. Scientific Reports [Internet]. 2016;6(June):1–12. Available from: https://www.nature.com/articles/srep26525.pdf

34. McCaig CD, Rajnicek AM, Song B, Zhao M. Controlling cell behavior electrically: current views and future potential. Physiological reviews. American Physiological Society; 2005;

35. Heinrich MA, Alert R, LaChance JM, Zajdel TJ, Košmrlj A, Cohen DJ. Size-dependent patterns of cell proliferation and migration in freely-expanding epithelia. Rosenblatt J, Stainier DY, Kabla A, editors. eLife [Internet]. eLife Sciences Publications, Ltd; 2020 Aug;9:e58945. Available from: https://doi.org/10.7554/eLife.58945

36. Schopf A, Boehler C, Asplund M. Analytical methods to determine electrochemical factors in electrotaxis setups and their implications for experimental design. Bioelectrochemistry. Elsevier; 2016;109:41–48.

37. Li M, Wang X, Rajagopalan P, Zhang L, Zhan S, Huang S, et al. Toward Controlled Electrical Stimulation for Wound Healing Based on a Precision Layered Skin Model. ACS Applied Bio Materials. ACS Publications; 2020;

38. Ream RA, Theriot JA, Somero GN. Influences of thermal acclimation and acute temperature change on the motility of epithelial wound-healing cells (keratocytes) of tropical, temperate and Antarctic fish. Journal of experimental biology. The Company of Biologists Ltd; 2003;206(24):4539–4551.

39. Zhao M. Electrical fields in wound healing—an overriding signal that directs cell migration. Seminars in cell & developmental biology. Elsevier; 2009. p. 674–682.

