## Supplementary material for "Come together: bioelectric healing-on-a-chip": Electronic Supplementary Information

#### Contents

1. Experimental Methods
2. Device Stability
3. Supplementary Figures (Figs S1, S2) and Video Captions (Videos S1, S2)
4. References

#### 1. Experimental Methods

##### Cell Culture

Primary keratinocytes were harvested from mice (courtesy of the Devenport Laboratory, Princeton University) and cultured in E-medium supplemented with 15% serum and 50 μM calcium. Cells were maintained at 37 °C under 5% CO2 and 95% relative humidity. Cells were split before reaching 70% confluence and passage number was kept below 30 for all experiments.

##### Device fabrication and assembly

This device assembly is a modified version of the SCHEEPDOG bioreactor from our prior work, and our methods are similar to those published previously (1). The chamber geometry was designed in vector graphics software (Affinity Designer) and then simulated in finite element software (COMSOL) to predict the field geometry. The stencil defining the pattern was cut from a 250 μm thick sheet of silicone rubber (Bisco HT-6240, Stockwell Elastomers) by a computer-controlled cutter (Cameo, Silhouette). This stencil was applied to a 10 cm tissue culture dish (Falcon) and formed the outline of the electro-stimulation zone. Fibronectin was adsorbed to the dish's surface to provide a matrix for cellular adhesion (protein was dissolved to 50 μg/mL in DI water, applied to the dish for 30 min at 37 °C, then rinsed three times with DI water). A second silicone stencil defined two 3x10 mm microwells, and 10 μL of a seeding solution of cells (singulated keratinocytes suspended in media at a density of $2.0 \times 10^6$ cells/ mL) was added to each well. Cells were left to settle for 6 hours in a humidified chamber at 37 °C. After this settling period, the dish was filled with 10 mL of media and incubated for 14 hours to form confluent monolayers.

The three acrylic pieces comprising the reusable device insert were laser milled (VLS 3.5, Universal Laser Systems) out of a 5.2 mm thick acrylic sheet. These individual layers of acrylic were stacked, clamped and solvent welded together with acrylic cement (SCIGRIP 4, SCIGRIP Asssembly Adhesives) and set for 24 h. The lid cap for the assembly was cut from a 3 mm thick acrylic sheet and a self-adhesive 1 mm-thick silicone sheet was adhered to one side to provide a better seal against the device insert. All components were sterilized by exposure to 5 min UV radiation in a cell culture hood just before assembly.

After the cell incubation period, the stencil was removed and the device was assembled immediately. To fabricate the integrated salt bridges, 4% w/v agarose was melted into phosphate buffered saline pH 7.4 on a hot plate. Once fully melted, the agarose was cast into the three slots in the acrylic device to serve as bridges. Once the agarose bridges had solidified, 5-10 mL of PBS was added to each of the three saline reservoirs associated with the agarose bridges. Then, the device was inserted into the tissue culture dish and clamped against the silicone stencil using four modified C-clamps (Humboldt Manufacturing Co.), taking care to avoid trapping air bubbles. Next, the chloridized silver electrodes were inserted into the lid cap and pressed against the acrylic device, so each electrode rested within a separate saline reservoir. Then, a Ti-wire recording electrode was inserted into each well, until it contacted one of the agarose bridges. The completed assembly was then moved to the microscope for imaging.

**Electrode Preparation and Characterization**

The silver chloride electrodes used in convergent wound healing assays were prepared by electroplating silver chloride onto silver foil electrodes. To prepare them, silver foil electrodes were immersed in 0.1 M KCl and poised at 3-5 V against a titanium wire counter electrode to a target current density of 1 $mA/cm^2$ and plated for 14-16 hours. Bleach immersion electrodes were prepared for the basis of comparison by submerging clean silver foil in a commercial 8% bleach solution for 24 hours.

We characterized the cathodic performance of these electrode materials by using a potentiostat to sweep five sequential cyclic voltammetry (CV) cycles in a three-cell electrochemical cell. The working electrode was 3 $cm^2$ of the cathode material under test, the counter electrode was a 15 $cm^2$ silver chloride foil, and the reference was a standard Ag/AgCl reference (SYC Technologies, Inc.). Normal operating conditions in our device show a cathodic potential of -1.5 to -1.7 V vs. Ag/AgCl when delivering a current of 6-8 mA, so voltage was swept from 0 to -2 V vs. Ag/AgCl.

**Instrumentation.**

Two Keithley source meters (Keithley 2400/2450, Tektronix) supplied current to the stimulation electrodes, and both sources shared the central cathode. A USB oscilloscope (Analog Discovery 2, Digilent Inc.) measured the chamber voltage at the Ti recording electodes, and both meters shared the central probe. A custom MATLAB script adjusted the output currents using proportional feedback control to maintain the desired field strength, and stimulation between the two pairs were alternated every 30 seconds so only one source was active at a given time.

**Microscopy**

All images were acquired on an automated Zeiss (Observer Z1) inverted fluorescence microscope equipped with an XY motorized stage controlled by Slidebook (Intelligent Imaging Innovations, 3i). The microscope was fully incubated at 37 °C within a polycarbonate enclosure. A peristaltic pump (Instech Laboratories) inside the chamber perfused fresh media through the electro-bioreactor at a rate of 2 mL/h. Polyolefin tubing with 1/32" inner diameter and 3/32" outer diameter (Flexelene™, United States Plastic Corp.) conducted media from the pump to the device. This tubing was connected to the acrylic device via a contact fit with a 2.0 mm inlet/outlet access holes cut in the device lid. To regulate media pH, 5% $CO^2$ was continuously bubbled through the media reservoir. All imaging used a 5X/0.16 fluorescence objective. Cells were either labeled by a lipophilic cytoplasmic dye (CellBrite Red, Biotium) and imaged with a Cy5 filter set ($\lambda_{ex}$/ $\lambda_{em}$ 644/665 nm) and 300 msec exposure per image or labeled with a live nuclear dye (NucBlue, Invitrogen) and imaged with a DAPI filter set ($\lambda_{ex}$/ $\lambda_{em}$ 358/461 nm) and 200 msec exposure per image. The outline of the central electrode slit was visualized by illumination with the DAPI filter set and a one-time 300 msec exposure. Fluorescence illumination was supplied by a metal halide lamp (xCite 120, EXFO). Images were captured at 10 min intervals.

**Image Processing**

Tissue velocity maps were generated using PIVLab, a MATLAB script performing FFT-based PIV (2). Iterative window analysis was performed using first 160×160 µm windows followed by 80×80 µm windows, both with 50% step overlap. Vector validation excluded vectors beyond five standard deviations and replaced them with interpolated vectors. Line integral convolution was used to visualize the flow field at certain time points and presented as the flow map. The velocity vector fields were then imported into MATLAB for kymograph generation and plotting. When required, labeled nuclei were tracked using FIJI's TrackMate plugin set to detect spots via Laplacian of the Gaussian filtering and to link using Linear Motion Tracking (3).

#### 2. Device Stability

##### Electrochemical stability

The results of electrochemical characterization of bare silver, bleached silver, and electroplated silver electrodes are shown in Fig. S1. In the case of bare silver, there is very little current until a reduction current caused by electrolytic evolution of $H_2$ from -1.5 V to -2.0 V, accompanied by visible formation of bubbles on the working electrode. Chloridized silver electrodes ideally would sink cathodic current by the breakdown of AgCl, thus avoiding this hydrolysis reaction. The bleached silver electrode had an expected linear CV for the first cycle, but after five cycles and just 0.7 mAh of charge transfer, the CV reverted to that of bare silver, as the chloridization layer has been consumed. The electroplated electrode shows a robust response that is nearly unchanged throughout 2.1 mAh of charge transfer throughout these five cycles. Because current delivered in the convergent field device under normal operation ranges from 6-8 mA, the equivalent current sunk by a cathode must be at 72-96 mAh for a 12 h experiment. That means that bleached silver electrodes are not sufficient for high, sustained current deliveries, and electroplated silver electrodes should be used instead, since a much larger AgCl volume can be plated onto the cathode than is formed by bleach immersion.

#### 3. Supplementary Figures and Video Captions

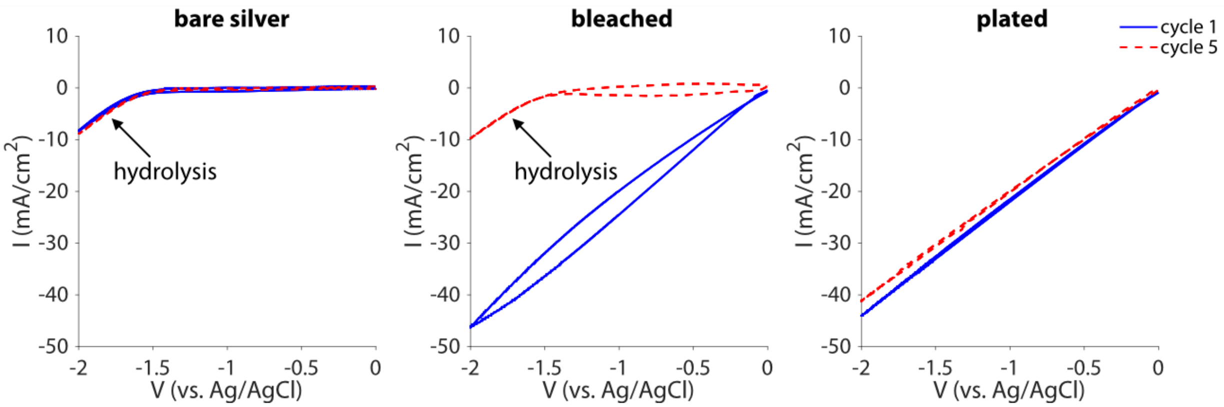


**Fig. S1. Electrochemical stability of three different cathode preparations, each probed by cyclic voltammetry.** Cyclic voltammograms measured by a potentiostat in a three-electrode system. The working electrode was made from the material under test, cut to an area of 3 $cm^2$. The counter electrode was a large silver chloride foil (15 $cm^2$ area), and the reference was a standard Ag/AgCl electrode. Five consecutive voltage sweeps were performed at a 100 mV/s scan rate from 0 to -2 V vs. Ag/AgCl. The first (solid blue line) and fifth (dashed red line) cycles are presented. The operating point of the cathode during standard device operation is -1.5 to -1.7 V vs. Ag/AgCl. At this voltage, bare silver produces faradaic current via hydrolytic evolution of hydrogen gas. Chloridizing the silver enables the electrode to source high faradaic current via the cathodic breakdown of AgCl at these voltages, avoiding potentially harmful pH shifts caused by hydrolysis. However, the method of chloridization matters, as bleach-immersed AgCl electrodes were stripped by the fifth cycle while electroplated AgCl electrodes lasted longer.

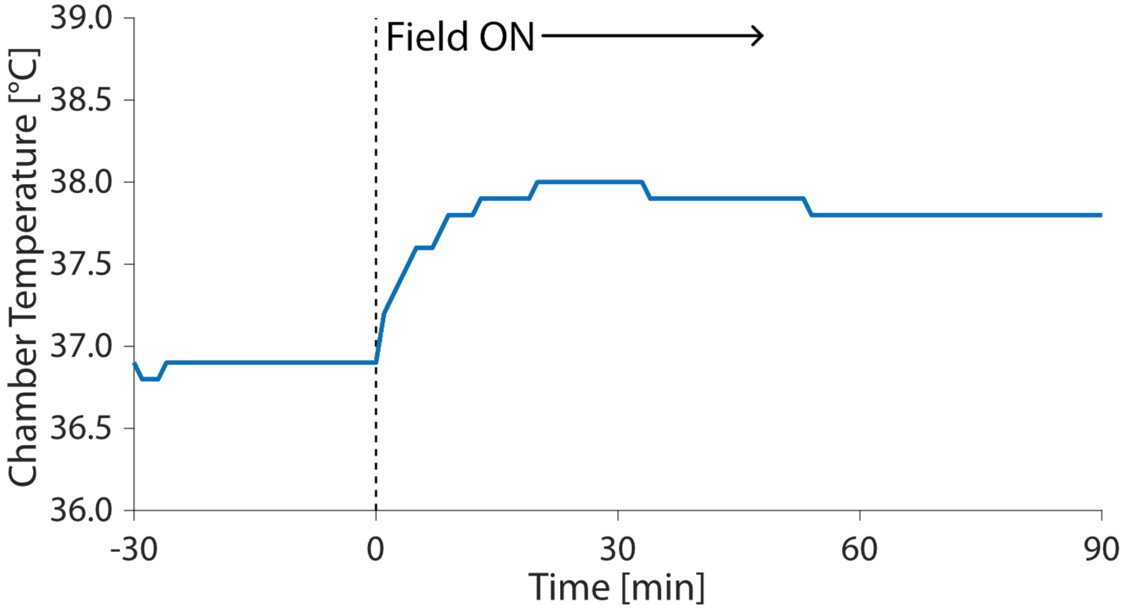


**Fig. S2. Temperature stability of device during stimulation.** Temperature measured by inserting a thermocouple into device's outlet port during perfusion. Stimulation field turned on at t = 0, marked by the dotted line. After the onset of stimulation, the system temperature reaches steady state in ~15 min, resulting in a temperature increase of ~1 °C due to Joule heating.

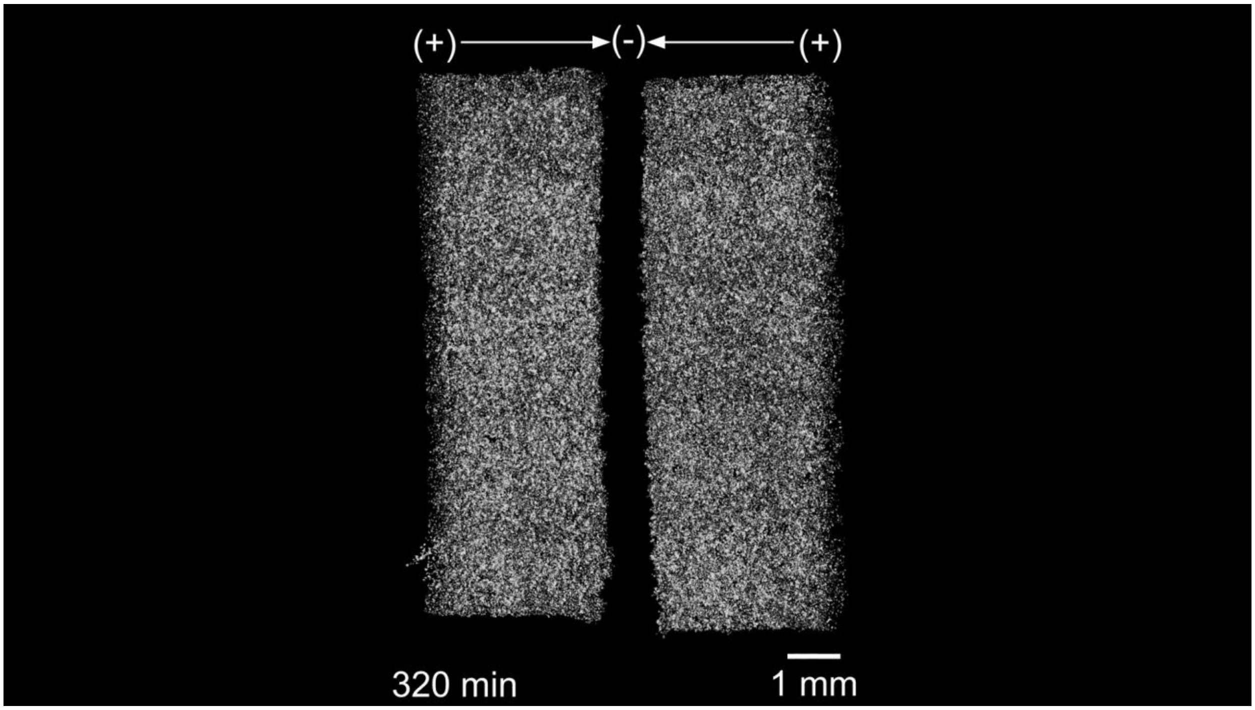


**Video S1: Time-lapse movie of converging stimulation of keratinocyte monolayers.** Following a 30-minute control period, a convergent field was applied to the center for 12 hours, alternating stimulation between the left and right tissues every 30 seconds. Cells were stained by a lipophilic Cy5 dye (imaged with $\lambda_{ex}$/ $\lambda_{em}$ 644/665 nm).

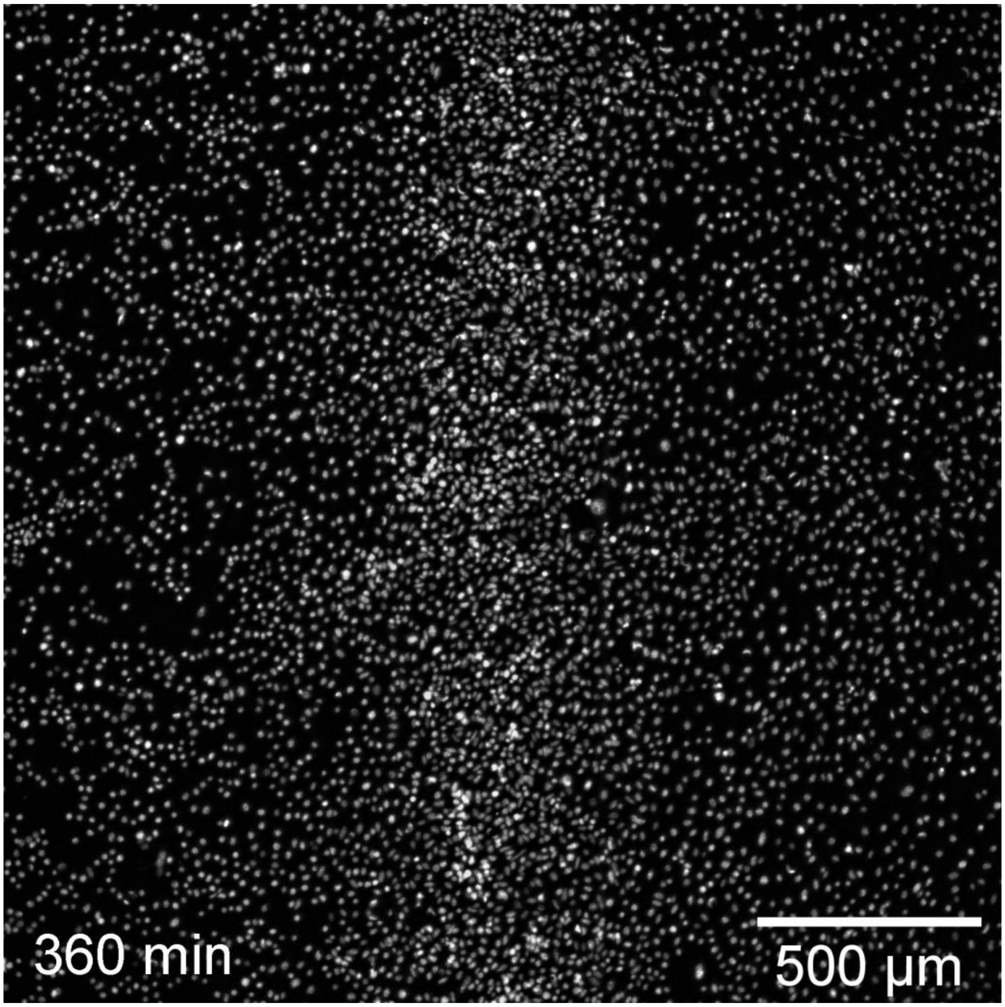


**Video S2: Time-lapse movie of convergence zone of a convergently stimulated keratinocyte monolayer.** A convergent field was applied to the center of a confluent monolayer for 6 hours, alternating stimulation between the left and right half every 30 seconds. Cells were stained by a live nuclear dye (imaged with $\lambda_{ex}$/ $\lambda_{em}$ 358/461 nm).

### 4. References


1. Zajdel TJ, Shim G, Wang L, Rossello-Martinez A, Cohen DJ. SCHEEPDOG: Programming Electric Cues to Dynamically Herd Large-Scale Cell Migration. Cell Systems [Internet]. Elsevier Inc.; 2020;10(6):506–514.e3. Available from: https://doi.org/10.1016/j.cels.2020.05.009
2. Thielicke W, Stamhuis E. PIVlab–towards user-friendly, affordable and accurate digital particle image velocimetry in MATLAB. Journal of open research software. Ubiquity Press; 2014;2(1).
3. Tinevez J-Y, Perry N, Schindelin J, Hoopes GM, Reynolds GD, Laplantine E, et al. TrackMate: An open and extensible platform for single-particle tracking. Methods. Elsevier; 2017;115:80–90.
